# Transcriptional and epigenomic profiling identifies YAP signaling as a key regulator of intestinal epithelium maturation

**DOI:** 10.1101/2023.05.10.540120

**Authors:** Laura M. Pikkupeura, Raul B. Bressan, Jordi Guiu, Yun Chen, Martti Maimets, Daniela Mayer, Pawel J. Schweiger, Stine L. Hansen, Grzegorz J. Maciag, Hjalte L. Larsen, Kadi Lõhmussaar, Marianne Terndrup Pedersen, Joji M. Yap Teves, Jette Bornholdt, Vladimir Benes, Albin Sandelin, Kim B. Jensen

## Abstract

During intestinal organogenesis, equipotent epithelial progenitors mature into phenotypically distinct stem cells that are responsible for life-long maintenance of the tissue. While the morphological changes associated with the transition are well-characterized, the molecular mechanisms underpinning the maturation process are not fully understood. Here, we leverage intestinal organoid cultures to profile transcriptional, chromatin accessibility, DNA methylation and 3D chromatin conformation landscapes defining fetal and adult epithelial cells. We observed prominent differences in gene expression and enhancer activity, accompanied by changes in 3D organization and local changes in DNA accessibility and methylation, between the two cellular states. Using integrative analyses, we identified sustained YAP transcriptional activity as a major gatekeeper of the immature fetal state. We found the YAP-associated transcriptional network to be regulated at various levels of chromatin organization, and likely to be coordinated by changes in extracellular matrix composition. Altogether, our work highlights the value of unbiased profiling of regulatory landscapes for the identification of key mechanisms underlying tissue maturation.

## Introduction

The fetal intestine undergoes vast expansion and remodeling, leading to the formation of rudimentary villi and a continuous intervillus space during development (Guiu and Jensen, 2015; Chin et al., 2017). Following villus formation, equipotent epithelial progenitors give rise to functionally defined adult stem cells (Barker et al., 2007; Guiu et al., 2019), which, following birth, become confined to the bottom of the intestinal crypts and are responsible for replenishment of the epithelium throughout life (Gehart and Clevers, 2019). Although the histological and morphological changes during this transition from the fetal to the adult intestine has been extensively described (Bjerknes and Cheng, 1981; Kim et al., 2007; Shyer et al., 2015; Shyer et al., 2013; Sumigray et al., 2018; Guiu et al. 2019; Maimetts et al., 2022), how the process is orchestrated at the molecular level remains largely unexplored.

Here, we harnessed organoid cultures to investigate cell-intrinsic molecular determinants of epithelium maturation. Analogous to their in vivo counterparts, stem cells derived from the adult intestinal epithelium give rise to budding organoids that contain both crypt-and villus-like domains, partially recapitulating the morphology and cellular composition of the adult epithelium (Sato et al. 2009). In contrast, fetal progenitors cultured under identical conditions form intestinal cystic spheroids that are mostly composed of undifferentiated progenitors and do not spontaneously mature in vitro. Interestingly, however, upon transplantation into the adult intestinal niche, fetal organoids can engraft and differentiate into mature cell types (Fordham et al. 2013; Mustata et al. 2013; Guiu et al. 2019), suggesting that their immature state in vitro is sustained by cell-intrinsic mechanisms.

Using this tractable model system, we employ a combination of RNA expression, chromatin accessibility, DNA methylation and 3D chromatin conformation profiling techniques to define transcriptional and regulatory landscapes that define the two developmental states. We find that fetal and adult epithelial cells are transcriptionally distinct, and that the activity of the regulatory elements is reflected by changes in local chromatin accessibility, DNA methylation, as well as chromatin interactions and compartmentalization. Furthermore, we identify changes in expression of extracellular matrix components and activation of YAP signaling as a candidate regulatory mechanism of the fetal to adult transition. Our work supports a model in which cell-intrinsic mechanisms contribute to the specification of adult stem cells and provide a framework for further exploration of determinants of tissue maturation.

## Results

### Fetal and adult intestinal organoids are transcriptionally distinct

Under growth-permissive conditions *in vitro*, intestinal adult stem cells self-organize to form budding organoids, whereas progenitors derived from the fetal epithelium grow as cystic spheroids (Fordham et al., 2013; Sato et al., 2009). Importantly, the two *in vitro* systems recapitulate the cellular composition of their i*n vivo* tissue counterparts, thereby providing a tractable platform to investigate cell intrinsic mechanisms without the confounding influences of microenvironmental or broader systemic cues. To characterize the differences between fetal progenitors and adult stem cells, we derived 3D cultures from the fetal (Embryonic day 16.5) and adult proximal part of the murine small intestine as described previously (Fordham et al., 2013; Mustata et al., 2013; Sato et al., 2009) - hereafter referred to as fetal enterospheres (FEnS) and adult organoids (aOrg), respectively. Established cultures were subjected to conditions that suppress differentiation (namely, EGF, Noggin, R-spondin1, CHIR99021 and Nicotinamide – ENR+ChNic) to minimize the influences of divergent cell type compositions between aOrg and FEnS (Figure 1A, S1A-B).

**Fig. 1.**
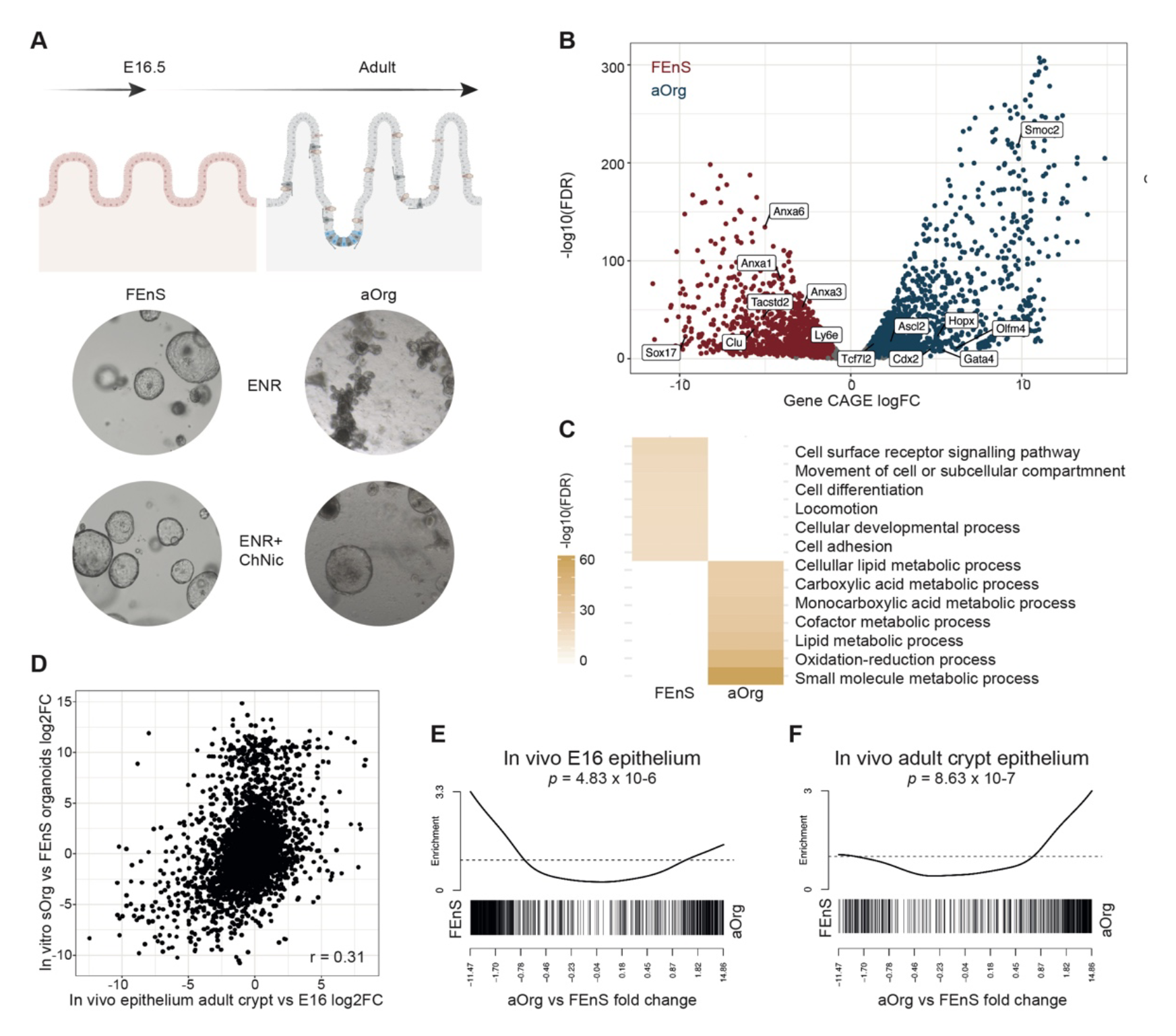
Fetal progenitors and adult stem cells are transcriptionally distinct and correspond to their *in vivo* counterparts. **(A)** Embryonic day 16.5 (E16.5) fetal and adult small intestinal epithelium derived organoids grown in ENR (Epidermal growth factor, Noggin, and R-Spondin1) and ENR+ ChNic (CHIR99021 and Nicotinamide). (**B)** Volcano plot of CAGE differential expression aggregated to gene-level. X axis shows adult vs. fetal log2 fold changes (FC), y axis shows log10 transformed *FDR* values. Colors: red = enriched in FEnS (*FDR* < 0.05 and log2FC < −1), blue = enriched in aOrg (FDR < 0.05 and log2FC > 1, and grey = not differentially expressed. Known state specific genes are labelled in the plot. (**C)** GO term enrichment among genes upregulated in FEnS and aOrg compared to all expressed genes. Rows indicate GO terms and columns FEnS or aOrg. Color indicates the significant level of enrichment (-log10 FDR). (**D)** Correlation plot of *in vivo* adult crypt vs E16 epithelium log2 fold changes and *in vitro* adult vs E16 derived organoids log2 fold changes. Pearson correlation coefficient is indicated with r. (**E)** Gene set enrichment analysis of the *in vivo* E16 genes in CAGE data sorted by aOrg vs FEnS fold change. (**F**) Gene set enrichment analysis of the *in vivo* adult crypt gene genes in CAGE data sorted by aOrg vs FEnS fold change.

We subsequently profiled the genome-wide activity of transcription start sites (TSSs) in the two systems using Cap Analysis of Gene Expression (CAGE), a technique that allows simultaneous profiling of messenger (mRNA) and enhancer RNA (eRNA) expression (Takahashi et al., 2012). Strikingly, based on mRNA levels, we found that 35% out of all expressed genes (11,386 genes) were differentially expressed between the two systems (*FDR* < 0.05 and absolute log2 fold change > 1; Figure 1B, S1E; Table S1). Significantly upregulated genes in adult state included known markers of adult intestinal stem cells (e.g *Olfm4*, *Ascl2*, *Cdx2*, *Gata4*, *Tcf7l2, Smoc2* and *Hopx*), while fetal-specific genes included *Sox17, Clu*, *Tacstd2, Anxa1, Anxa3* and *Anxa6* (Figure 1B). Gene ontology (GO) analysis showed that the mRNAs upregulated in FEnS were enriched in cell surface receptor signaling pathways, locomotion, development and adhesion (Figure 1C). As previously described (Yui et al., 2018), several of these over-represented terms were linked to a set of FEnS-upregulated extracellular matrix genes. Conversely, the genes upregulated in the aOrg were mostly enriched in GO-terms associated with metabolic processes (Figure 1C). Importantly, comparison to the transcriptome of freshly isolated epithelial cells matching these developmental stages (Maimets et al. 2022) indicated significant conservation of fetal-and adult-specific signatures in the established cultures. The *in vivo* E16.5 fetal vs adult crypt log2 fold changes correlated significantly with i*n vitro* aOrg vs FEnS log2 fold changes (Pearson r = 0.31, *p* < 2.2e-16; Figure 1D). Furthermore, the in vivo fetal and adult specific gene sets were enriched among the FEnS and aOrg genes, respectively (Figure 1E-F). Altogether this indicates that, despite identical culture conditions and morphological similarities, 3D cultures derived from fetal and adult epithelium display distinct transcriptional programs that reflect their original developmental state.

### Fetal and adult states have discrete enhancer and promoter landscapes

To explore the mechanism underlying the transcriptional differences, we next set out to characterize the enhancer and promoter landscapes in FEnS and aOrg by combining the CAGE data with ATAC-seq profiling of chromatin accessibility. Consistent with (Andersson et al., 2014b), the combined analysis of the datasets showed that 90% of CAGE reads fell within ATAC-seq peaks (Figure S2A). By intersecting open chromatin ATAC-peaks with active eRNA or mRNA CAGE peaks, we defined 13,064 candidate enhancers and 13,325 active gene promoters (Figure 2A). As expected, ENCODE ChIP-seq transcription factor peaks were enriched at the enhancer candidate regions, when compared to ATAC-seq peaks without CAGE signal (Figure S2B; Fisher’s exact test *p* < 2.2×10^-16^, odds ratio 3.15) suggesting that these are functional elements. Moreover, we found that 1,700 (13%) out of 11,386 genes had one or more alternative promoters (Figure S2C). Differential expression analysis of annotated promoters between FEnS and aOrg revealed that 2,271 (17%) out of 13,361 active promoters were significantly upregulated in the FEnS and 2,440 (18%) in the aOrg (*FDR* < 0.05 and absolute log2 fold change > 1; Figure 2B). Similarly, 3,240 (23%) of the identified enhancer candidates were upregulated in the FEnS state in terms of eRNA expression (*FDR* < 0.05 and absolute log2 fold change > 1) and 4,063 (29%) in the aOrg (Figure 2B). Since eRNA expression has been shown to correlate with enhancer activity (Andersson et al., 2014a), these observations suggest that the state-specific activity of regulatory elements underlies the vast gene-expression differences between the fetal and adult states.

**Fig. 2.**
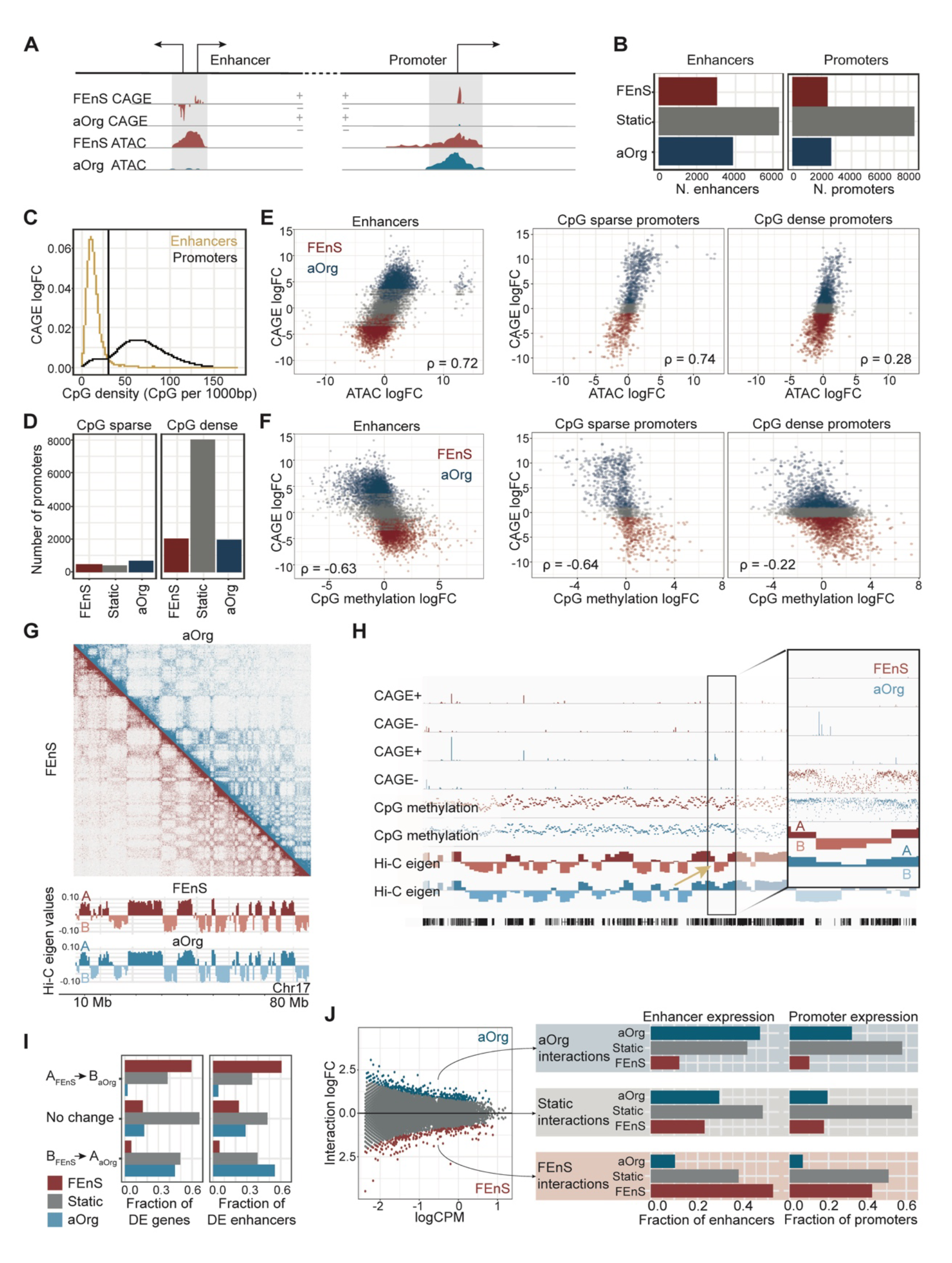
Chromatin changes in different levels reflect changes in transcription. **(A)** Illustration of typical enhancer and promoter, defined as transcribed ATAC-seq peaks. (**B)** Number of FEnS (red), static (grey), and aOrg (blue) enhancers (left) and promoters (right). (**C)** CpG density distribution in enhancers (yellow) and promoters (black) within 1kb windows from enhancer center or promoter summit, respectively. (**D)** Number of differentially expressed promoters among CpG sparse promoters (< 30 CpGs per 1kb; left) and CpG dense promoters (right). (**E)** Correlation between ATAC logFC and CAGE logFC at enhancers and CpG sparse and dense promoters. Each dot corresponds to one predicted enhancer or gene promoter, colored by differential expression by CAGE. Spearman’s correlation coefficient is indicated with *ρ*. (**F)** Correlation between CpG methylation logFC and CAGE logFC at enhancers and CpG sparse and dense promoters. Each dot corresponds to one predicted enhancer or gene promoter, colored by differential expression by CAGE. Spearman’s correlation coefficient is indicated with *ρ*. (**G)** Top: A representative Hi-C chromatin interaction heatmap showing chr17 in 500kb bins. X and Y axis correspond to chr17 coordinates. Colors indicate interactions between chromosomal regions in FEnS (red) and aOrg (blue) small intestinal epithelium derived organoids. Bottom: a bar plot showing corresponding eigenvalues of the first principal component of 500kb bins along chromosome 17 for each state. Both plots are based on averages of two biological replicates. The color represents the state (FEnS = red & aOrg = blue) and shade represents compartment (A = dark, B = light). (**H)** Tracks of chromosome 19 showing CAGE signal from + and - strand, CpG methylation % per CpG, eigenvalues of first principal component of 500kb Hi-C bins and NCBI RefSeq mm10 gene annotations. Zoom-in contains a B_FEnS_ -> A_aOrg_ transitioning region. The yellow arrow points to the region transitioning between the compartments in the zoom-in. (**I**) Bar plot shows the fraction of genes and predicted enhancers significantly more active in FEnS (red), aOrg (blue), or not significantly changing between states (grey), located in regions constitutively located in the same compartment or moving from A_FEnS_ -> B_aOrg_ or B_FEnS_ -> A_aOrg_ compartments. (**J)** Right: differential interactions intensities between 100kb bins shown as an MA plot with mean interaction counts per million (CPM) on X axis and interaction log2 fold changes (FC) on Y axis. Interaction, where intensities change significantly between the states (differential interactions, FDR < 0.05) are colored by the direction of the change (blue – aOrg; red – FEnS: grey – static interactions). Left: bar plots show number of enhancers (left) and promoters (right) within interacting bins, split by whether interactions are static or enriched in fetal or adult or static. Bar colors represent the differential expression status of the enhancers and promoter by CAGE (FDR < 0.05 and absolute log2FC > 1); X axis indicates fraction of differentially expressed enhancers and promoters overlapping the differentially interacting bins. Y axis indicates the direction of differential expression.

### State-specific enhancers are enriched in clusters

Given the observed changes in enhancer accessibility and the reported role of enhancer clusters in driving cell type-specific transcription (Hnisz et al., 2013; Whyte et al., 2013), we next set out to assess the overall enhancer distribution in each state. Interestingly, approximately 50% of the identified enhancers were identified in proximity (12,500 bp) to at least one other enhancer candidate, forming 2,211 enhancer clusters (Figure S2C; Table S2). Enhancers within larger clusters have been reported to be co-regulated (Hah et al., 2015; Schmidt et al., 2015). Indeed, 1,046 enhancer clusters showed high eRNA expression correlation (Figure S2D) between associated enhancers. FEnS-and aOrg-specific enhancers were enriched within these internally correlated clusters compared to single enhancers or enhancers in uncorrelated clusters (Fisher’s exact test *p* < 2.2×10^-16^, odds ratio 3.75; Figure S2E). This raises the possibility that internally correlated enhancer clusters may underpin the regulation of the distinct cell states.

### Chromatin accessibility and CpG methylation changes reflect transcriptional changes at enhancers and a subset of promoters

We hypothesized that state specific enhancer and promoter activity would reflect differences in chromatin accessibility. Enhancers and promoters differ in their GC content, which affects chromatin structure. In particular, promoter nucleosome binding has been shown to correlate with GC density, with GC dense regions being inherently nucleosome repelling (Fenouil 2012, Saxonov 2006). We observed a bimodal distribution of promoter GC density with a smaller peak of GC sparse promoters with a density comparable to enhancers, which tend to have low GC density (Figure 2C). Based on this observation, we divided promoters into “CpG sparse” and “CpG dense” and strikingly the majority of CpG sparse promoters were differentially expressed between the FEnS and aOrg, while this was not the case for CpG dense promoters (Figure 2D). To explore this further, we assessed the correlation of chromatin accessibility (ATAC) and their change of expression (CAGE). The changes in chromatin accessibility of CpG sparse promoters strongly correlated with gene expression (Spearman *ρ* = 0.74), similar to enhancers. In contrast, the CpG dense promoters exhibited only weak correlation (Spearman *ρ* = 0.28) and chromatin remained accessible irrespective of expression status (Figure 2E, S2F). Specifically, 22% of FEnS-and 29% of adult-specific enhancers exhibited differences in the levels of activation in the respective state (FDR < 0.05 and absolute log fold change > 1). Similarly, 22% of FEnS-and 29% aOrg-specific CpG sparse promoters were differentially accessible, while this was only the case for 8% of FEnS-and 8% of aOrg-specific CpG rich promoters. We conclude that chromatin accessibility at enhancer regions and a subset of promoters with low CpG density is dynamically regulated across developmental states and co-occurs with expression changes. In contrast, the majority of promoters are constitutively accessible despite prominent transcriptional changes.

Given the correlation between dynamic chromatin accessibility and promoter GC content, we speculated that enhancer and promoter activity at these GC sparse regions would correlate with DNA methylation. We, therefore, analyzed differences in CpG cytosine methylation between FEnS and aOrg cultures using whole genome bisulfite sequencing. Indeed, we observed a strong negative correlation between expression and methylation at enhancers (Spearman *ρ* = - 0.63) and CpG sparse promoters (Spearman *ρ* = −0.64), but only a weak negative correlation at CpG rich promoters (Spearman *ρ* = −0.22), which were constitutively hypomethylated (Figure 2F, S2F). In conclusion, we found that chromatin accessibility and DNA methylation that are associated with patterns of eRNA expression define each developmental stage. In contrast only a few promoters showed state-specific chromatin changes, with a vast majority of them being constitutively accessible and hypomethylated despite the changes in gene expression. Altogether, this suggests that rather than by local chromatin changes at promoters, developmental programs may be largely regulated by differential enhancer usage.

### Large-scale chromatin structure is broadly conserved in fetal and adult organoids

Since we hypothesized that the developmental gene expression programs are regulated by differential enhancer usage, rather than local changes at promoters, we next enquired whether changes in the 3D chromatin structure linking enhancers and promoters would be an important determinant of state-specific transcription. Given the evidence for the role of the genome compartmentalization into active (A) and inactive (B) chromatin compartments for cell fate transitions (Lieberman-Aiden et al. 2009; Dixon et al. 2015; Hu et al. 2018; Stadhouders et al. 2019; Johnstone et al. 2020), we first combined *in situ* DNase Hi-C with DNA methylation profiling to investigate these large-scale chromatin changes. Using an eigenvector-based approach on 500kb genomic bins, we defined A and B compartments, and observed that the overall chromatin architecture was largely maintained between the two states (Figure 2G). Importantly, the Hi-C eigenvalues were negatively correlated with published data of interactions between chromatin and nuclear lamina (Meuleman et al. 2013), with negative eigenvalues (B compartment) corresponding to regions with strong lamina interactions (Figure S2G). This is consistent with the notion that nuclear lamina interactions are key for the spatial organization of chromatin, wherein B compartment regions associate closely with lamina associated domains (LADs), and that most LADs are constitutively associated with the nuclear lamina regardless of cell type (Meuleman et al. 2013; van Steensel and Belmont 2017).

Interestingly, we noted that although the overall chromatin structure was largely unchanged (Figure 2G), some regions switched compartments between fetal and adult cells. Specifically, 73 regions moved from active to inactive compartment (A FEnS → B aOrg), and 81 regions had the reciprocal pattern (B FEnS → A aOrg), while the remaining 4,787 bins were static (either A or B in both FEnS and aOrg; Figure S2H). To investigate whether the compartment switches reflected the levels of chromatin accessibility, CpG methylation and transcriptional activity, we overlaid the Hi-C data with ATAC, DNA methylation and CAGE expression datasets. We observed that these few large compartment transitions were accompanied by changes in CpG methylation levels and transcriptional activity (Figure 2H). In general, CpG methylation levels were higher in A compartment and lower in B compartment and compartment transitions coincided with corresponding changes in methylation levels (Supplementary Figure S2I). Consistent with previous observations (Berman et al. 2011; Fortin and Hansen 2015; Johnstone et al. 2020), we noticed that broad regions with low levels of methylation and disordered methylation patterns preferentially overlapped with the B compartment (Supplemental Figure S2J-K), confirming the compartment annotation based on the Hi-C data.

We next explored whether these few changes in chromatin compartmentalization correlated with activation of state-specific enhancer landscapes and gene regulatory networks. Indeed, we found that A FEnS → B aOrg transitioning regions were enriched for FEnS-upregulated genes and enhancer candidates, whereas aOrg-upregulated genes and enhancer candidates were enriched in B FEnS → A aOrg regions (Figure 2I)

### State-specific changes in chromatin interactions reflect transcriptional changes

Despite the few compartmental changes, differential interaction analysis at smaller scale (100kb resolution), identified 3,226 regions with significantly different interaction intensities in FEnS and aOrg (FDR < 0.05; Figure 2J), which directly correlated with promoter and enhancer expression changes between the two states (Figure 2J). Altogether, this indicates that, despite conservation of the large-scale chromatin structure, smaller-scale chromatin reorganization correlates with transcriptional activity driven by specific regulatory elements in the fetal and adult states.

Finally, to further investigate the fine-scale changes in chromatin organization, we defined changing chromatin boundaries corresponding to changes in topologically associated domain structure (n=3,113; Figure S2L-M). Intriguingly, enhancers that occurred in clusters were generally in closer proximity (< 50kb) to these changing chromatin boundaries when compared to single enhancers (Fisher’s exact test *p* = 6.9×10^-7^, odds ratio 1.26). In addition, we found that differentially expressed genes were enriched in regions adjacent (< 50kb) to changing chromatin boundaries (Fisher’s exact test *p* = 1.1×10^-5^, odds ratio 1.22; Table S3). Collectively, these analyses demonstrate that local chromatin rearrangements might underlie state-specific differential enhancer and transcription activity between fetal and adult cells.

### Different transcription factors are associated with stage-specific chromatin changes

Given the reported role of lineage-specific transcription factors (TFs) as major determinants for cellular identity, we reasoned that the differences in gene expression are likely to be driven by stage-specific TFs. Indeed, we found that 159 and 144 TFs were significantly upregulated in FEnS and aOrg, respectively (FDR < 0.05 and absolute log fold change > 1; Figure 3A). These included TFs previously described to play roles in cellular identity in the fetal and adult intestine, such as Sox17 and Gata4, respectively (Figure 3A-C) (San Roman et al., 2015; Miura & Suzuki, 2017). Importantly, stage-specific enhancers and promoters were enriched for predicted binding sites of the differentially expressed TFs. FEnS promoters and enhancers were enriched for motifs associated with AP1, TEAD, SOX and NF-kB TF families, while aOrg enhancers and promoters showed enrichment of predicted TF binding sites (TFBSs) for HNF4A, CDX2 and GATA4 (Figure 3D). Furthermore, the observed TFBS enrichment was more prominent at enhancers than at promoters. This is consistent with the observation that TFs exert their role predominantly at enhancer regions (Whyte et al., 2013) and suggests that fetal progenitors and adult stem cells are regulated by different gene regulatory networks that activate divergent enhancer landscapes.

**Fig. 3.**
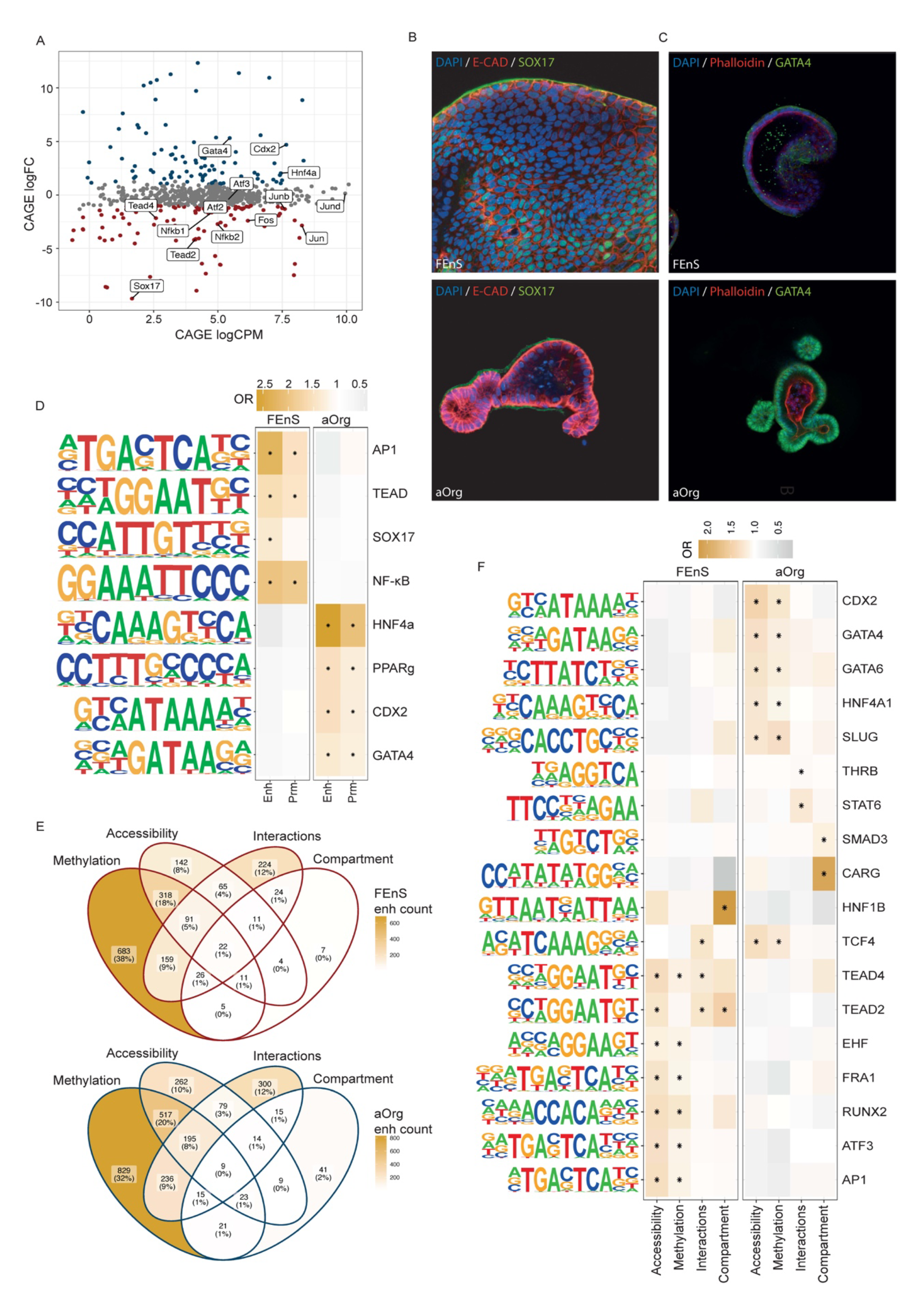
Distinct transcription factor networks drive fetal and adult state specific promoter and enhancer activity. (**A**) MA plot of transcription factor gene level aggregated CAGE expression log2 counts per million (CPM) over log2 fold changes (FC). The transcription factors with motifs enrichment (as in Figure 3D) in FEnS promoters and enhancers are labelled red, and the factors with motifs enrichment in aOrg promoters and enhancers are labelled blue. (**B**) Representative immunofluorescence images showing DAPI (blue), E-Cadherin (red), and SOX17 (green) (**C)** DAPI (blue), Phalloidin (red), and GATA4 (green) in FEnS and aOrg. (**D)** Transcription factor binding site enrichment at enhancers and promoters. Each row represents one transcription factor or a group of transcription factors sharing a binding motif. Color represents enrichment odds ratios over all expressed enhancers and promoters, respectively. Enrichment with FDR value < 0.05 are marked by asterisks (*). **(E)** Venn diagram depicting overlap of differential methylation (WGBS), accessibility (ATAC), interactions (Hi-C), and compartment status (Hi-C) among FEnS (top) and aOrg (bottom) enhancers. (**F)** Transcription factor binding site enrichment at differentially methylated, accessible, interacting, or compartment switching enhancers compared to all state-specific enhancers. Each row represents one transcription factor or a group of transcription factors sharing a binding motif. Color represents enrichment odds ratios over all expressed enhancers and promoters, respectively. Enrichment with FDR value < 0.05 are marked by asterisks (*).

We next used TFBS enrichment analyses to explore whether TF activity might regulate activity of state-specific enhancers that display changes in either methylation, accessibility, 3D interactions or compartmentalization status. We found that different TFBSs were enriched at enhancers that were either differentially methylated (WGBS) or accessible (ATAC) or had differential 3D chromatin interactions or moved between chromatin compartments, when compared to the remaining FEnS-or aOrg-specific enhancers. This was consistent with the observation that changes in accessibility most often co-occur with changes in methylation, but less frequently with changes in interactions or compartmentalization (Figure 3E). Importantly, regardless of the mode of regulation, FEnS and aOrg enhancers showed enrichment for different classes of TFBSs (Figure 3F). Within a developmental state, the same TFBSs were enriched in differentially methylated and accessible enhancers, which differed from the TFBSs enriched at differentially interacting or compartment switching enhancers located within A FEnS → B aOrg or B FEnS → A aOrg regions (Figure 3F). Interestingly, TEAD (TEAD2 and TEAD4) motifs were consistently enriched at both the locally differentially accessible and methylated FEnS enhancers, as well as at enhancers associated with local interaction and compartment changes, suggesting that TEAD-mediated transcription might be a major determinant of the fetal-specific transcriptional program.

### Fetal progenitors are maintained by sustainably high YAP activity levels

Given the consistent occurrence of predicted binding sites for TEAD and AP1 factors, known co-effectors of YAP signaling (Zanconato et al., 2015; Torato et al., 2018), in both local and larger-scale chromatin changes, we hypothesized that YAP transcriptional activity might be involved in the maintenance of the fetal progenitor state. In agreement, we found that FEnS are highly enriched for the canonical YAP transcriptional signature (Gregoriff et al. 2015) and express high levels of the YAP direct target genes *Ankrd1*, *Ctgf*, *Cyr61* and *Clu* (Figure 4A- B). In agreement, image analysis confirmed high levels of nuclear (active) YAP in FEnS, compared to fewer cells - mostly restricted to the bud domains - with nuclear signal in aOrg (Figure 4C). Of note, using a published dataset (Maimets et al., 2022), we found that the YAP-associated transcriptional signature is also enriched in the fetal epithelium compared to adult crypt cells *in vivo* (Figure 4D-E). Akin to the *in vitro* cultures, nuclear YAP signal was also observed throughout the E16.5 epithelium, but only at the upper part of adult crypts (Figure 4F). Altogether, this indicates that maintenance of fetal epithelium both *in vitro* and *in vivo* is supported by high YAP activity.

**Fig. 4.**
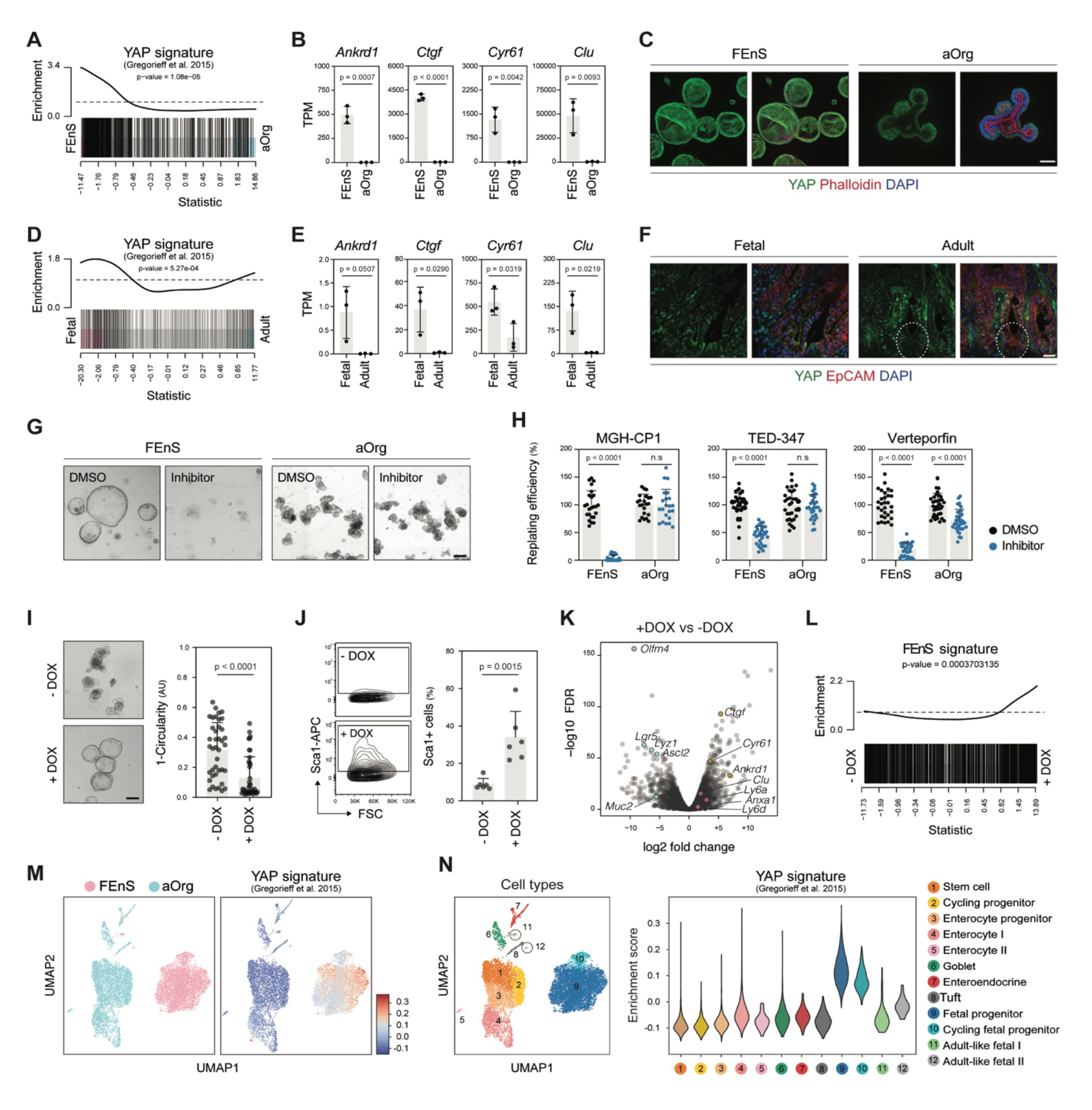
Fetal cells are maintained by sustained high levels of YAP activity. **(A)** GSEA showing enrichment of YAP-associated gene signature in FEnS relative to aOrg cultures. **(B)** Bar plots depicting expression levels of direct YAP target genes in FEnS and aOrg cultures. **(C)** Immunofluorescent analysis showing YAP subcellular localization in FEnS and aOrg cells. Nuclear signal is observed in the majority of fetal cells, while adult cells show mostly cytoplasmic localisation. Scale bar: 100µm. **(D)** GSEA showing enrichment of YAP-associated signature in E16.5 epithelial progenitors versus adult crypt cells *in vivo*. **(E)** Bar plots depicting expression levels of direct YAP target genes in E16.5 epithelial progenitors and adult crypt cells. **(F)** Immunofluorescent analysis showing YAP subcellular localization in E16.5 epithelial progenitors and adult crypts cells in vivo. Scale bar: 20µm. **(G)** Representative phase contrast images of FEnS and aOrg cultures treated with YAP signaling inhibitors. Images from MGH-CP1 treatments are shown. Scale bar: 200µm. **(H)** Bar plots depicting replating efficiency following treatment with the indicated inhibitors. Each dot represents a technical replicate of at least three independent experiments. **(I)** Left - Representative images of TetON-YAP aOrg cultures in the absence and presence of doxycycline (-/+DOX). Right - Bar plot depicting quantification of organoid circularity. Scale bar: 200µm. **(J)** Analysis of Sca1 protein expression by flow cytometry in tetON-YAP aOrg cultures. **(K)** Volcano plot showing differentially expressed genes upon DOX treatment. Representative adult-and fetal-specific genes are highlighted in blue and red, respectively. Known YAP target genes are highlighted in yellow. **(L)** GSEA showing enrichment of FEnS-associated gene signature upon DOX treatment. **(M)** Left - UMAP visualization of single cells from FEnS and aOrg cultures colored by sample. Right - Average expression of YAP signature genes overlaid on the UMAP plot cell type. **(N)** Left **-** UMAP visualization of single cells from FEnS and aOrg cultures colored by cell type. Clusters were annotated based on expression of known marker genes. Right - Enrichment analysis of a YAP signature separated by each cluster. See associated manuscript for method details.

To test the requirement for YAP activity, we next treated fetal and adult derived organoid cultures with three specific pharmacological inhibitors of YAP-TEAD mediated transcription: MGH-CP1, TED-347 and Verteporfin (Li et al., 2020; Bum-Erdene et al., 2019; Liu-Chittenden et al., 2012). Using replating assays, we consistently found that organoids derived from fetal state are more sensitive to transient inhibition of the pathway, indicating that growth of fetal cells is more dependent on YAP activity than adult cells (Figure 4G-H).

To test whether high YAP activity was sufficient to induce a fetal-like state, we next derived adult cultures from an inducible TetON-hYAP/H2B-mCherry mouse strain (see methods), in which addition of doxycycline (DOX) induces expression of a constitutively active (S127A mutant) YAP transgene. Using this system, we found that YAP induction following organoid passaging causes aOrg to grow as cystic, fetal-like round structures, with concomitant upregulation of the fetal-specific cell surface marker Sca1 (Yui et al., 2018; Figure 4I-J). Transcriptionally, YAP overexpression led to the downregulation of a number of stem cell markers and a global upregulation of the fetal-specific gene signature (Figure 4K-L). Interestingly, however, DOX-treated aOrg cultures did not fully recapitulate fetal cultures transcriptionally, nor could these mutant aOrg be maintained long-term in the normal medium conditions in the presence of DOX (data not shown), indicating that these are not fully converted into a self-sustained fetal state.

Finally, using single cell RNA-seq profiling of fetal and adult organoids, we found that a small fraction of the cells in the fetal cultures spontaneously mature into adult–like secretory cell types (Figure 4M; Hansen and Larsen et al., submitted). Interestingly, while YAP-associated genes were highly expressed in the majority of fetal cells, these ‘maturing’ cells (cluster 11 and 12) showed a low enrichment in the signature, suggesting that the maturation process entails a reduction in YAP activity (Figure 4M-N). Altogether, these data indicate that activated YAP is necessary for the fetal state and sufficient to induce a fetal-like transcriptional program.

### Differential YAP activity levels correlate to differential expression of ECM genes

Given the role of YAP as a mechanoresponsive factor (Panciera et al., 2017), we speculated that differential expression of extracellular matrix (ECM) components might underlie the differential activation levels observed in fetal and adult cells. In agreement, we found fetal-specific genes to be highly enriched in genes associated with cell adhesion and cell surface receptors (Figure 1C). Similarly, extracellular matrix related GO terms were also enriched among genes regulated at different levels (Figure S3). Among these genes, we found various laminin subunits, collagens and fibronectin to be expressed at higher levels in the fetal cells, both *in vitro* and *in vivo* (Figure S4A).

As signaling by various ECM components have been shown to trigger YAP activation through focal adhesion kinase (FAK) and Src family kinases (Panciera et al., 2017), we tested the effects of pharmacological inhibitors of the kinases in culture. Consistent with the previous results using YAP inhibitors, fetal cultures showed increased sensitivity to FAK and SRC inhibitors, confirming the requirement of the pathways and active YAP signaling for maintenance of the fetal cells (Figure A4B-C)

## Discussion

Cellular state transitions such as the maturation of fetal progenitors into the adult tissue stem cells require activation of specific transcriptional programs. In the intestinal epithelium, recent works indicate that maturation is associated with enhancer decommissioning, changes in chromatin accessibility as well as differential requirements for mitochondrial activity (Banerjee et al., 2018; Chen et al., 2019; Srivillibhuthur et al., 2018, Jadhav et al., 2019). Here, we used organoid cultures to further provide a comprehensive characterization of transcriptional and chromatin landscapes of fetal progenitors and adult stem cells of the intestinal epithelium. Our analyses indicate that, despite few changes in chromatin compartment and promoter accessibility, widespread differences in gene expression and enhancer usage characterize each developmental stage. Together with TF motif analyses, these observations suggest that maturation is regulated by vastly different enhancer landscapes associated with distinct transcription factor networks. Importantly, we found that a considerable fraction of state-specific enhancers was organized in clusters, a feature previously suggested to drive expression of genes important for cellular identity (Hnisz et al., 2013; Whyte et al., 2013).

Integrative analysis of state-specific regulatory elements and chromatin features identified TEAD and AP1 proteins - transcriptional cofactors for YAP signaling – as potential regulators of the fetal transcriptional program. Given the well described role of YAP signaling in fetal development and tissue repair (Azzolin et al. 2014; Yui et al. 2018; Zheng and Pan 2019), we hypothesized that sustained YAP activity is a key player in maintenance of the fetal state. Indeed, we showed that fetal cultures cells are more dependent on YAP activity and the upstream activating signals mediated by FAK and Src. In line with the elevated expression ECM proteins, we speculate that the deposition of such components by fetal progenitors represents a cell-intrinsic mechanisms to sustain YAP activity and, ultimately, reinforce the progenitor state and/or prevent their premature transition into functionally restricted adult intestinal stem cells. *In vivo*, such a mechanism is likely to operate in synergy with changes in the stroma and niche formation (Guiu and Jensen, 2015; Gehart and Clevers, 2019) during morphogenesis to ensure proper timing of intestinal stem cell maturation.

Experimentally, the identification of such cell-intrinsic mechanisms might assist the improvement of ES/iPS cell differentiation protocols *in vitro*. It is evident from numerous studies that the derivation of tissue specific cell types from pluripotent stem cells is complicated by an inability to direct the cells into an adult state (Gieseck et al., 2014; Kroon et al., 2008; Parent et al., 2013; Sun et al., 2013; Trounson and Dewitt, 2012; Woodford and Zandstra, 2012). Although this maturation process has been achieved upon transplantation and exposure to complex *in vivo* environments in some cases (Cohen and Melton, 2011; Parent et al., 2013; Sun et al., 2013; Watson et al., 2014), the development of more robust differentiation protocols would require the identification of signaling pathways and underlying cell-intrinsic mechanisms.

In summary, we demonstrate that fetal and adult intestinal progenitor/stem cells are maintained by distinct gene regulatory networks, mostly operating on the enhancer landscape. We propose that such networks provide an additional mechanism to safeguard the successful maturation and/or differentiation of the epithelium during development. Similar mechanisms are likely to operate in the context of tissue repair and cancer, given the emerging evidence that these processes often require reacquisition of developmental transcriptional programs (Yui et al., 2018; Vasquez et al., 2022). Indeed, we and others have previously found that regeneration of the adult intestinal epithelium entails a YAP-mediated reprogramming process into a fetal-like state (Yui et al., 2018; Ayyaz et al., 2019; Martin-Alonso et al., 2021). We, therefore, envisage that the comprehensive dataset presented here will provide a framework for further investigation of determinants of fetal progenitor and adult stem cell states, and shed light on conserved mechanisms of tissue repair and neoplasia. For instance, the two adult-specific regulators *Cdx2* and *Gata4* (Verzi et al., 2010) are similarly surrounded by adult state specific enhancer clusters and have been implicated in intestinal regeneration and/or tumorigenesis. As such, we generated a user-friendly webpage that allows visualization of differentially expressed promoters and enhancers, as well as DNA methylation and chromatin structure patterns (see data availability statement for details).

## Materials and Methods

### Mice

C57BL/6J mice (Taconic) were used for all the experiments, unless otherwise specified. Transgenic. TetON-hYAP/H2B-mCherry was generated through breeding of a Tet-YAP strain (Jansson and Larsson 2012) with a Tet-H2B-mCherry strain (Jackson Laboratory #014602). None of the animals had been subjected to prior procedures. All animals were housed in specific-pathogen-free animal facilities under a 12-h light–dark cycle always with companion mice. Food and water were provided *ad libitum*. The Danish Animal Inspectorate has approved all animal procedures.

### Adult and fetal epithelium derived organoid cultures

Organoids were grown as previously described (Fordham et al., 2013; Yui et al. 2018). Briefly, epithelial fragments from fetal intestine (intestines from three fetuses were pooled for each sample) and scraped adult crypts (from one individual per sample) were harvested from the proximal parts of the small intestine using EDTA (2mM). Harvested epithelial fragments adult crypts were embedded in Matrigel droplets and cultured in advanced DMEM/F12 with GlutaMAX and Penicillin/Streptomycin supplemented with EGF (Peprotech; 50 ng/ml), Noggin (Peprotech; 100ng/mL), and R-spondin1 (R&D Systems; 500 ng/ml). When indicated, the growth medium was supplemented with CHIR99021 (Stemgent; 3μM) and Nicotinamide (Sigma; 10 mM). For transgene induction and the TetON-YAP-H2B-mCherry, doxycycline (Sigma #D9891, 1ug/ml) was added for 48h immediately after passaging.

### Replating assays

FEnS and aOrg cultures were passaged using conventional mechanical disruption, and 72h after incubating with the pharmacological inhibitors for 24h. DMSO was used as control. Treated organoids were passaged again and incubated with control media. Organoid formation efficiency (relative to DMSO controls) was determined 72h after. The following pharmacological inhibitors were used: MGH-CP1 (Selleckchem #S9735, 20uM), TED-347 (Selleckchem #S8951, 20uM), Verteporfin (Sigma #SML0534, 0.25uM), Dasatinib (Sigma #SML2589, 0.25-1uM) and PF573228 (Tocris #3239, 2.5-10uM).

### CAGE

CAGE libraries for three biological replicates of fetal spheroids and adult organoids were prepared as previously described (Takahashi et al., 2012) with an input of 3000 ng of total RNA. All primers and adapters were purchased from Integrated DNA Technologies (IDT). After individually preparing CAGE libraries for each sample, three samples with unique barcodes (CTT, GAT, and ACG) were pooled per sequencing lane and 30% Phi-X was spiked into each lane. Sequencing was carried out at the National High-throughput DNA sequencing Centre, University of Copenhagen, with Illumina HiSeq2000.

### CAGE data processing

CAGE data was processed as previously described (Boyd et al., 2018). Briefly, reads with identical barcodes were matched to their originating samples. After removal of linker sequences and filtering, reads were mapped to the mm10 assembly with Bowtie2 (Langmead et al., 2009) and only uniquely mapping reads not mapping to chrM were retained for further analysis. 5’ ends of CAGE tags (CTSSs) supported by only a single read were excluded from the analysis. CTSSs were subsequently Tags Per Million (TPM) normalized (CAGE reads per total mapped reads in library times 10^6^). CTSSs on the same strand within 20bp of each other were merged into tag clusters (TCs) and quantified in all samples (single read CTSSs were included for the quantification). A summit was identified in each TC, defined as the single base-pair position within the TC with highest total TPM coverage across all samples.

### ATAC-seq data processing

Paired-end reads were mapped with Bowtie2 (Langmead, Trapnell, Pop, & Salzberg, 2009) against mm10 assembly and only uniquely mapping reads were retained using samtools (Li et al., 2009). Subsequently, duplicate reads were removed and regions highly contaminated by duplicate reads (99.9th percentile) as well as ENCODE black list regions were masked out (Amemiya, Kundaje, & Boyle, 2019). Reads aligning to plus strand were offset by +4 bp and reads aligning to minus strand with −5bp as described in (Buenrostro et al., 2013). The peaks were called for each replicate using MACS2 without a shifting model and the background lambda as local lambda. Differential accessibility analysis was performed separately for promoters and enhancers using generalized linear model quasi-likelihood framework of edgeR (Robinson et al., 2010).

### Identification of promoters and enhancers and gene-level expression

As we found that only 10% of CAGE signals fell outside of ATAC-seq peaks, only CTSSs overlapping an ATAC-seq peak were retained. Using NCBI RefSeq mm10 annotation, we identified peaks overlapping transcription start sites (+/- 500bp) or exons (+/- 200bp) as gene promoters and intronic and intergenic peaks as potential enhancers. For promoters, we retained ATAC peaks overlapping a TC > 0.8 average TPM in all three replicates in at least one condition. For enhancers, a noise cutoff was estimated for the sum of the CAGE TCs overlapping an ATAC peak for all unique ATAC peak lengths. For gene level expression, the sum of TCs overlapping the gene on the same strand was used. To identify enhancer clusters, we stitched non-exon overlapping transcribed ATAC peaks outside within 12.5 kb from each. Among these clusters, we calculated the pairwise Pearson correlations between all member enhancers and the clusters with average pairwise correlation > 0.5 were defined as internally correlated enhancer clusters.

### Exploratory CAGE analysis and differential expression

Principal component analysis was performed for combined TPM normalized CAGE counts at promoter and enhancers using prcomp function from base R with scaling and centering. Differential expression analysis was performed separately at gene, promoter, and enhancer levels using generalized linear model quasi-likelihood framework of edgeR (Robinson et al., 2010). For identifying a set of differentially expressed genes an absolute log2 fold change threshold of one was used and p-values were corrected for multiple testing using Benjamini-Hochberg method and FDR < 0.05 cut-off was used. For analysis of gene ontology term enrichment, topGO with Fisher’s exact test was used (Alexa A.; Rahnenfuhrer J, 2019). All expressed genes were used as a background set.

### Whole genome bisulfite sequencing

Genomic DNA of fetal and adult organoids (3 biologically independent replicates each) was purified using Genomic DNA Purification Kit (Thermo scientific). All 6 libraries were prepared with ACCEL-NGS METHYL-SEQ DNA Library Kit.

### Whole genome bisulfite sequencing data processing

After quality trimming with Trim Galore the reads were mapped and methylation status was determined with Bismark, which bisulfite converts the reads in silico prior to mapping with Bowtie2 to a bisulfite converted reference genome. (Krueger & Andrews, 2011). We found that the mapping efficiency was low due to the large number of chimeric reads. To mitigate this, the mapping was performed in two steps: first all reads were mapped in paired end mode with the option --unmapped, following which the unmapped R1 were mapped in single-end mode (directional mode) and the unmapped R2 in single-end mode (--pbat mode). The paired-end and single-end alignments were merged after methylation extraction.

### Identification of partially methylated domains (PMDs) and differentially methylated regions (DMRs)

To segregate the genome into PMDs and non-PMDs, we use MethylSeekR (Burger, Gaidatzis, Schubeler, & Stadler, 2013). We identified a consensus set of PMDs present in all 6 samples. Fetal and adult stage specific PMDs were defined as regions present in all replicates in one condition and none of the replicates in the other condition. We identified differentially methylated regions by BSmooth implemented in bsseq (Hansen, Langmead, & Irizarry, 2012). We used the methylation calls processed with Bismark and performed the smoothing and differential methylation analysis with BSmooth. We used only CpGs which were covered by more than four reads in all six samples. The smoothing was done for regions containing at least 70 CpG or were at least 2kb wide, whichever is largest, using the whole chromosomes as the smoothing cluster. For computing t-statistics, the variability was estimated based on the fetal samples. For identifying DMRs we used a cutoff 4.6, based on t-statistic quantiles.

### Hi-C

We performed *in situ* DNase Hi-C as previously described (Ramani et al., 2016). Briefly, 48 wells of fetal spheroids and adult organoids per replicate (2 biological replicates per condition), corresponding to approximately one million cells were released from Matrigel and fragmented by repeated washing with cold PBS with 0.1% BSA. The organoid fragments were cross-linked in 2% formaldehyde for 20 minutes, quenched, and washed. Following, the cells were lysed, and chromatin was digested with 2 units of DNase I for 5 minutes. The nuclei were purified with AMPure XP beads and washed followed by end repair and dA-tailing reactions. Biotin-labeled bridge adapters were then ligated overnight to the fragments followed by another round of purification with AMPure XP beads and washing. The nuclei were treated with PNK to phosphorylate adapters and ligation prior to reversal of cross-linking, DNA precipitation and purification. The ligation products with biotin were then pulled down with MyOne C1 beads followed by washing and end repair, dA-tailing, and ligation of sequencing adapters. The libraries were amplified for 14 PCR cycles with primers containing Nextera barcodes for sample multiplexing: N701 (TCGCCTTA), N702 (CTAGTACG), N703 (TTCTGCCT), and N704 (GCTCAGGA). Libraries were then purified and quantified with Qubit and Fragment Analyzer (Agilent) and the four libraries were pooled and paired-end sequenced with Illumina NextSeq.

### Hi-C data processing

Hi-C reads were matched to their originating samples. The reads were subsequently quality trimmed with Trim Galore. The reads were mapped and processed with HiC-Pro (Servant et al., 2015). We used MAPQ score threshold 30 and excluded read pairs mapping within 1kb, to retain valid pairs and generate ICE normalized contact maps at 100kb resolution. All valid read pairs were converted into Juicebox viewable.hic matrixes for visual exploration of interaction patterns (Durand et al., 2016) with KR (Knight-Ruiz) normalization (Knight & Ruiz, 2012) at 500kb resolution.

### Compartment analysis

A/B compartments were identified as previously described (Lieberman-Aiden et al., 2009). Briefly, we used Juicer to compute the Pearson’s correlation matrix of the observed/expected interaction matrix of contiguous non-overlapping 500kb bins and its first principal component (eigenvector) (Durand, Shamim, et al., 2016). The signs of eigenvectors were flipped according to gene density and the bins with positive eigenvalues were assigned as A and negative as B compartment. For stringency we excluded the bins where the eigenvector had an absolute value less than 0.01. We defined compartment changing regions as bins where the sign of the eigenvector was positive in both replicates of one condition and negative in other and magnitude larger than 0.01 in all samples.

### Correlation of eigenvalues with nuclear lamina interaction values

Hi-C eigenvalues were correlated with the average of previously reported genome-wide DamID lamin interaction values reported as log2-transformed Dam-fusion / Dam-only ratio of three different murine cell lines: neural precursor cells (NPCs), astrocytes (ACs), and embryonic fibroblasts (MEFs) (Meuleman et al., 2013).

### Differential interaction analysis

For the differential interaction analysis, we used the diffHic package (Lun and Smyth, 2015; Robinson et al., 2010). The Hi-C read pairs were summarized into pairs of contiguous non-overlapping 100kb bins, and bin overlapping repeat regions or pairs supported by fewer than ten read pairs were excluded from further analysis. We used the median abundance of inter-chromosomal bin pairs to estimate non-specific ligation rate and required over 5-fold higher abundance than this threshold, combined with a trended filter considering the interaction distance and required higher abundance than a fitted value for that distance, since larger counts are expected for shorter interaction distances. Furthermore, we excluded diagonal interactions within the same bin. After filtering the data was LOESS normalized, and significant differences were found using the generalized linear model quasi-likelihood framework and the generated P-values were corrected for multiple testing using Benjamini-Hochberg method.

### Boundary analysis

To identify differential domain boundaries, we used an approach based on directionality index implemented in diffHic (Dixon et al., 2015; Lun and Smyth, 2015). Briefly, by counting the difference in number of read pairs between the target 100kb bin and upstream or downstream 1Mb interval, a directionality index was calculated for each 100kb bin. After removing low-abundance bins, dispersion estimation and fitting of generalized linear model, bins where directionality statistic changes between fetal and adult states were identified after multiple testing correction with Benjamini-Hochberg method. We then identified a set of promoters within 50kb from a differential boundary and analyzed gene ontology term enrichment, using topGO with Fisher’s exact test. (Alexa A.; Rahnenfuhrer J, 2019).

### Transcription factor binding motif analysis

Transcription factor binding analysis was performed separately for enhancers and promoters. Enhancer regions were defined as +/- 300bp from enhancer midpoint and promoter regions as −500bp to +100bp around the CAGE summit. Differentially expressed enhancers and promoters in FEnS and aOrg were compared to the background set composed of all expressed enhancers or promoters, respectively. Next, we used the HOMER findMotifsGenome tool to calculate enrichment of previously determined motifs (Heinz et al., 2010). HOMER motif logos were extracted from http://homer.ucsd.edu/homer/motif/.

### RNA-seq

RNA was collected using RNAeasy Microkit (Qiagen). Libraries were prepared using TruSeq RNA kit (Illumina) following manufacturer instructions. The samples were sequenced with Illumina NextSeq. Reads were aligned using qAlign function from Bioconductor package QuasR (Gaidatzis et al, 2015) with default parameters. Mapped reads were assigned to known UCSC genes and quantified with qCount function from QuasR.

### RNA-seq exploratory analysis and differential expression

Differential expression analysis was performed using the generalized linear model quasi-likelihood framework of edgeR (Robinson et al., 2010). For identifying a set of differentially expressed genes P-values were corrected for multiple testing using Benjamini-Hochberg method and FDR < 0.05 cut-off was used. Gene set enrichment was performed with edgeR fry function for a list of YAP activated genes (Gregorieff et al., 2015), or the identified FEnS specific genes.

### Immunostainings

Fetal and adult organoids were plated in 96-well imaging plates (Greiner #655090) and fixed in PFA for 10 min at room temperature (RT). Cells were permeabilized and blocked in PBS-Triton X-100 0.2%, 2% normal goat serum, 1% BSA, 100 µM Glycine 100 µM for 60min at RT. Primary antibodies were diluted in PBS-TX100 0.2% + BSA 0.5% + NGS 1% and incubated overnight at 4 °C with slow agitation. Following washes with 0.1% Tween20 PBS, the appropriate Alexa Fluor-conjugated secondary antibodies (1:500) were incubated as the primary antibodies. Cells were counterstained with DAPI and Phalloidin for 20 min at RT, and RapiClear (Sunjin Lab #RC152001) solution was added before imaging.

For staining tissue samples, proximal halves of fetal and adult small intestines were collected, embedded in OCT and snap-frozen in dry ice. Tissue sections (7 μm thick) were prepared with a microtome, and subsequently were dried overnight at −80°C followed by 20 min at RT. Dried sections were fixed with ice-cold 4% PFA for 10 minutes. Blocking and permeabilization was performed in PBS containing 0.2% Triton X-100, 2% normal goat serum, 1% BSA and 100 µM Glycine. Primary antibodies were incubated overnight at 4°C in PBS containing 0.2% Triton X-100, 0.5% BSA and 1% normal goat serum followed by incubation with secondary antibodies in the same buffer for 90 min. All washes were performed with PBS containing 0.1% Tween-20. DAPI was used for nuclei staining. Images were acquired using laser scanning confocal microscope (Leica Stellaris 8) and analyzed with Fiji. The following primary antibodies were used: YAP (Cell Signaling Technologies #14074, 1:100), E-Cadherin (BD Biosciences #610281, 1:500), Sox17 (R&D systems #AF1924, 1:300) and Gata4 (Santa Cruz #sc-25310, 1:500).

### Flow Cytometry

Organoids were collected in 0.1% BSA solution, spun down and dissociated into single cells through incubation with TriplE solution at 37 degrees for 5 min. Following centrifugation, cells were resuspended in 1% BSA solution and incubated with Sca1-APC antibody (eBioscience #15-5981-83, 1:1000) for 15 min at RT. DAPI was used as viability staining. Data was collected with LSR Fortesa X20 (BD Bioscience) and analyzed with FlowJo.

## Acknowledgments

We thank members of the KBJ and AS labs for technical assistance and comments on the manuscript.

## Author contributions

Conceptualization - LMP, RBB, JG, AS and KBJ. Methodology - LMP, RBB, JG, YC, AS, and KBJ.; Investigation - LMP, RBB, JG, MM, PJS, SLH, HLL, KL and JB. Formal analysis - LMP, RBB, YC, DM and GJM. Resources - MM, MTP and SLH. Writing: Original Draft - LMP and RBB. Writing: Review & Editing - LMP, RBB, AS and KBJ. Supervision - AS and KBJ. Funding Acquisition - AS and KBJ.

## Data and materials availability

All raw and processed sequencing data generated in this study have been submitted to the NCBI Gene Expression Omnibus (GEO; https://www.ncbi.nlm.nih.gov/geo/) under accession numbers GSE228519. *In vivo* fetal data is available under accession number GSE183671. To facilitate the access to the community, we generated a user-friendly webpage that allows visualization of differentially expressed promoters and enhancers, as well as DNA methylation and chromatin structure patterns. The page can be accessed through the link: https://shiny.binf.ku.dk/fetal_adult_regulatory_app/

---

**Fig. S1.**
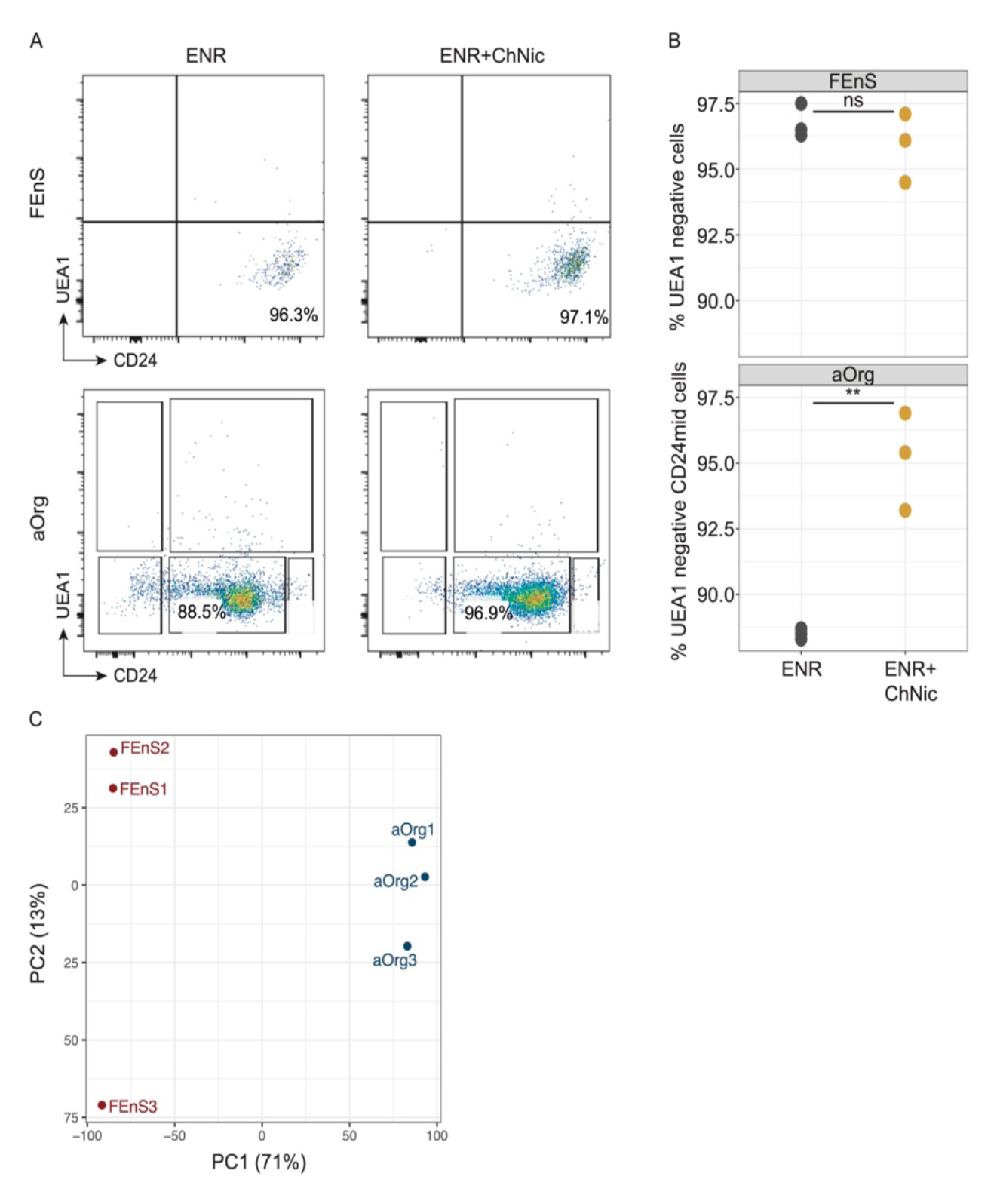
**(A)** Representative flow cytometry plots of CD24 surface expression vs Ulex Europaeus Agglutinin I (UEA1) binding in fetal spheroids and adult organoids grown in EGF, Noggin, and R-Spondin1 (ENR) (left) and ENR-CHIR99021-Nicotinamide (ENR-Ch-Nic) (right). Columns are ENR and ENR-Ch-Nic and rows FEnS and aOrg. (**B.** Quantification of UEA1 negative and positive cells based on flow cytometry analysis. Each dot represents a biological replicate. ns = non-significant based on two-sided t-test. **: *p* value < 0.01 based on two-sided t-test. (**C)** Principal component analysis based on CAGE expression. Shade indicates replicate and color indicates condition (Red – fetal; blue – adult).

**Fig. S2.**
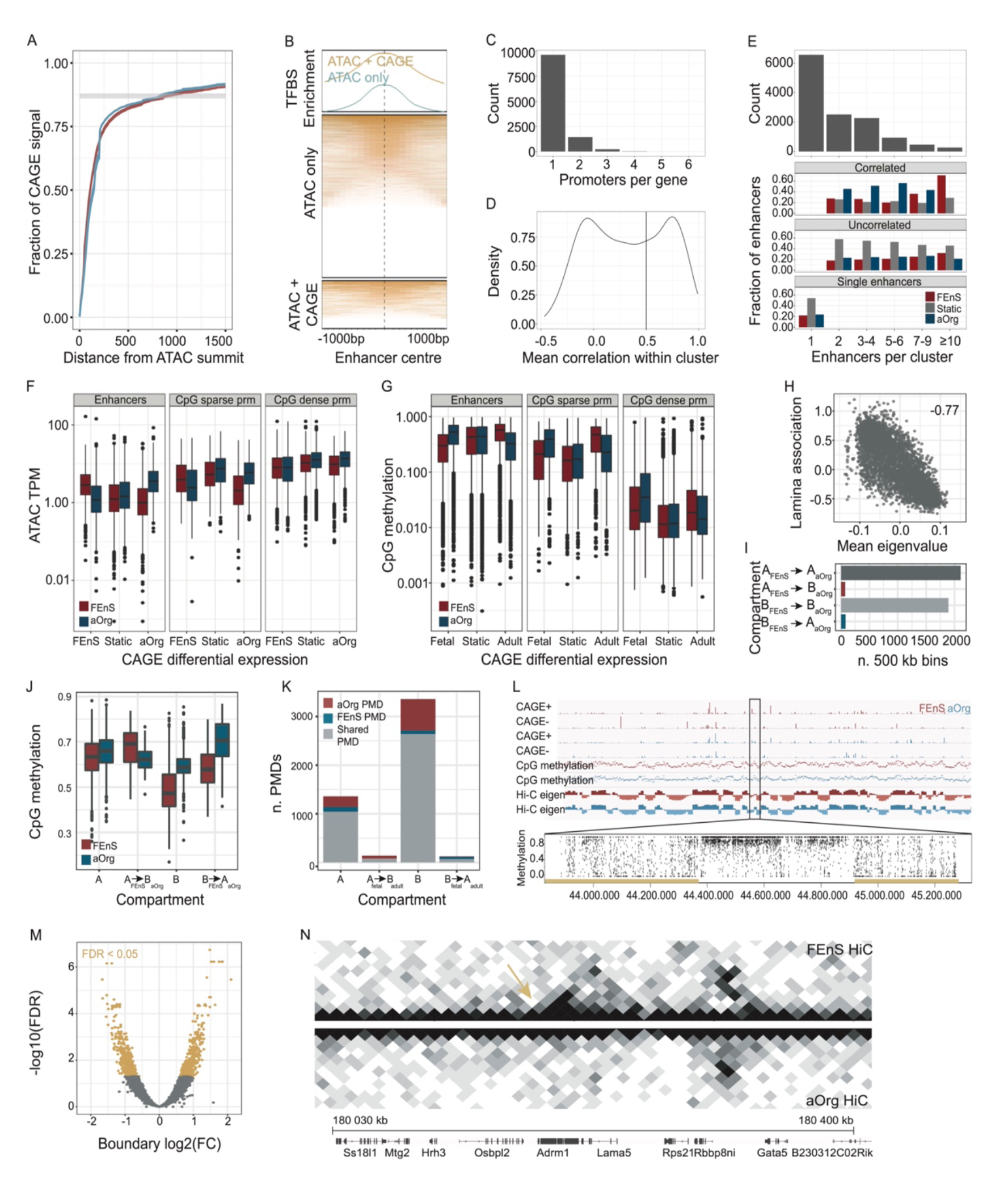
**(A**) Fraction of CAGE signal within area around ATAC-seq summits. (**B)** Heatmap illustrating enrichment of ENCODE transcription factor peaks within enhancer candidate regions defined based on ATAC-seq peaks (ATAC only) or transcribed ATAC-seq peaks (ATAC + CAGE). A density plot summarizes transcription factor peak enrichment on top of the heatmap. (**C)** Number of promoters per gene. (**D**) Distribution of average pairwise correlations between enhancers in clusters. (**E)** Number of enhancers located in clusters with different number of members (top) and fraction of differentially expressed enhancers among singleton enhancers and uncorrelated-and correlated enhancer clusters split by cluster size colored by differential expression. (**F)** Boxplot of normalized ATAC counts on Y axis at differentially expressed and static (based on CAGE data) enhancers (left) and promoters (right) shown as mean across two replicates of FEnS and aOrg samples. Color indicates condition (Red – FEnS; blue – aOrg). (**G)** Box plot of normalized average CpG methylation level on Y axis at differentially expressed and static (based on CAGE data) enhancers (left) and promoters (right) shown as mean across two replicates of fetal and adult samples. Color indicates condition (Red – FEnS; blue – aOrg). (**H)** Scatter plot shows average eigenvalues of adult and fetal HiC data vs. average genome-nuclear lamina interaction strength. Each dot corresponds to one 500kb region. Pearson correlation coefficient - 0.77, p < 2×10^-16^ (two-sided correlation test). (**I)** Bar plot shows the number of 500kb bins in all constitutive A and B compartments and regions moving between compartments. (**J)** Box plots of CpG methylation levels in A_fetal_ & A_adult_, A_FEnS_ -> B_aORg_, B_FEnS_ & B_aOrg_, B_fFEnS_ -> A_aOrg_ regions in FEnS and aOrg, indicated by color. The data represents averages over three biological replicates per condition. (**K)** Bar plot shows the number of shared and condition specific partially methylated domains (PMDs) in A_FEnS_ & A_aOrg_, A_FEnS_ -> B_aOrg_, B_FEnS_ & B_aOrg_, and B_FEnS_ -> A_aOrg_ regions. (**L)** Tracks of chromosome 18 organized as in Figure 2H. The zoom-in illustrates representative methylation patterns in A and B compartments. Partially methylated domains (PMDs) overlapping B compartment are labelled in yellow. **(M)** Volcano plot showing changes directionality statistic between FEnS and aOrg. 100kb bins with significant (FDR < 0.05) change in boundary strength between the states are colored yellow. (**N)** Example of FEnS specific boundary.

**Fig. S3.**
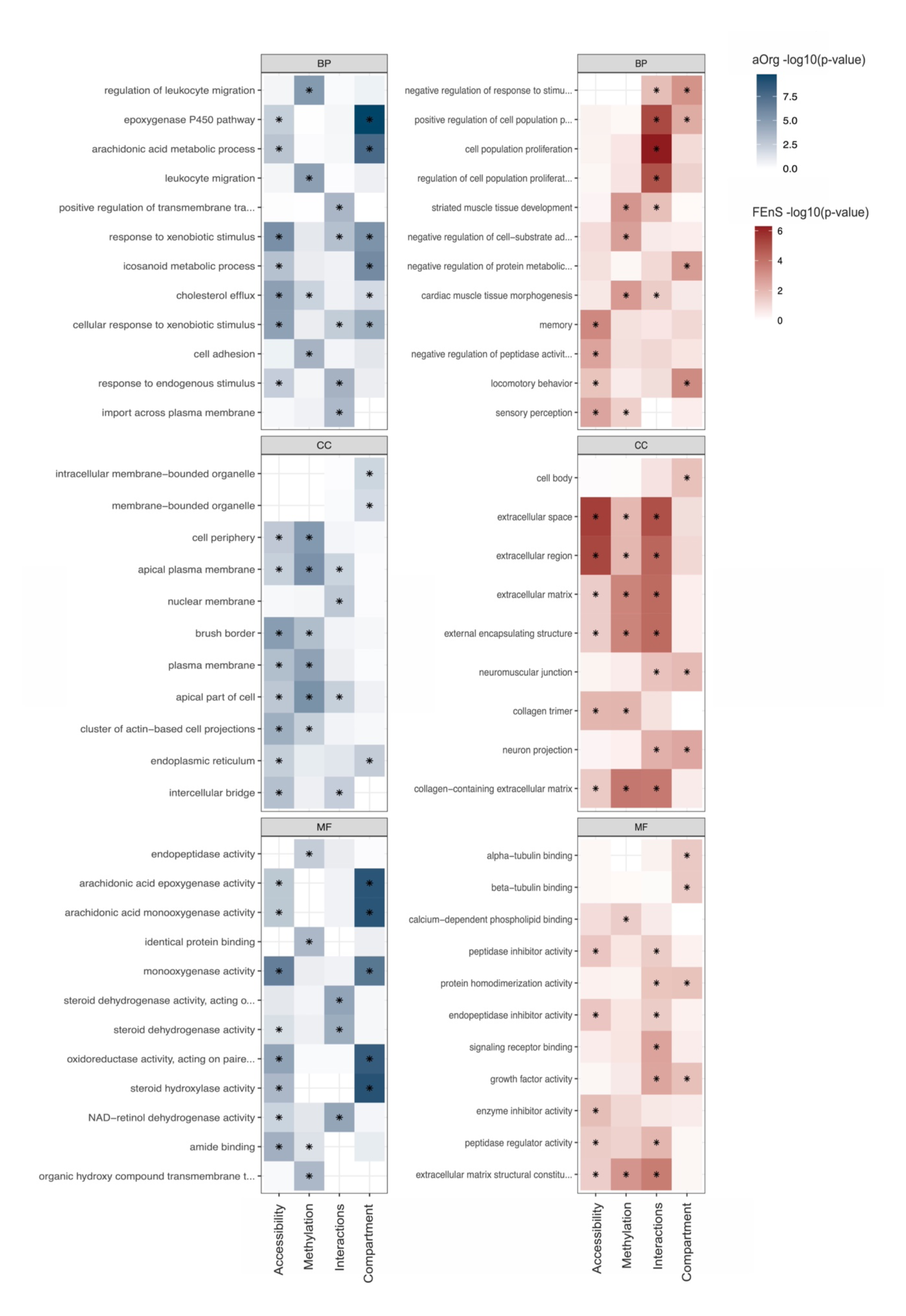
GO term enrichment among genes with differentially methylated, accessible, interacting, or compartment switching promoters in FEnS and aOrg compared to all FEnS and aOrg specific genes. Rows indicate GO terms and columns differentially methylated, accessible, interacting, or compartment switching promoters. Color indicates the level of enrichment (-log10 FDR).

**Fig. S4.**
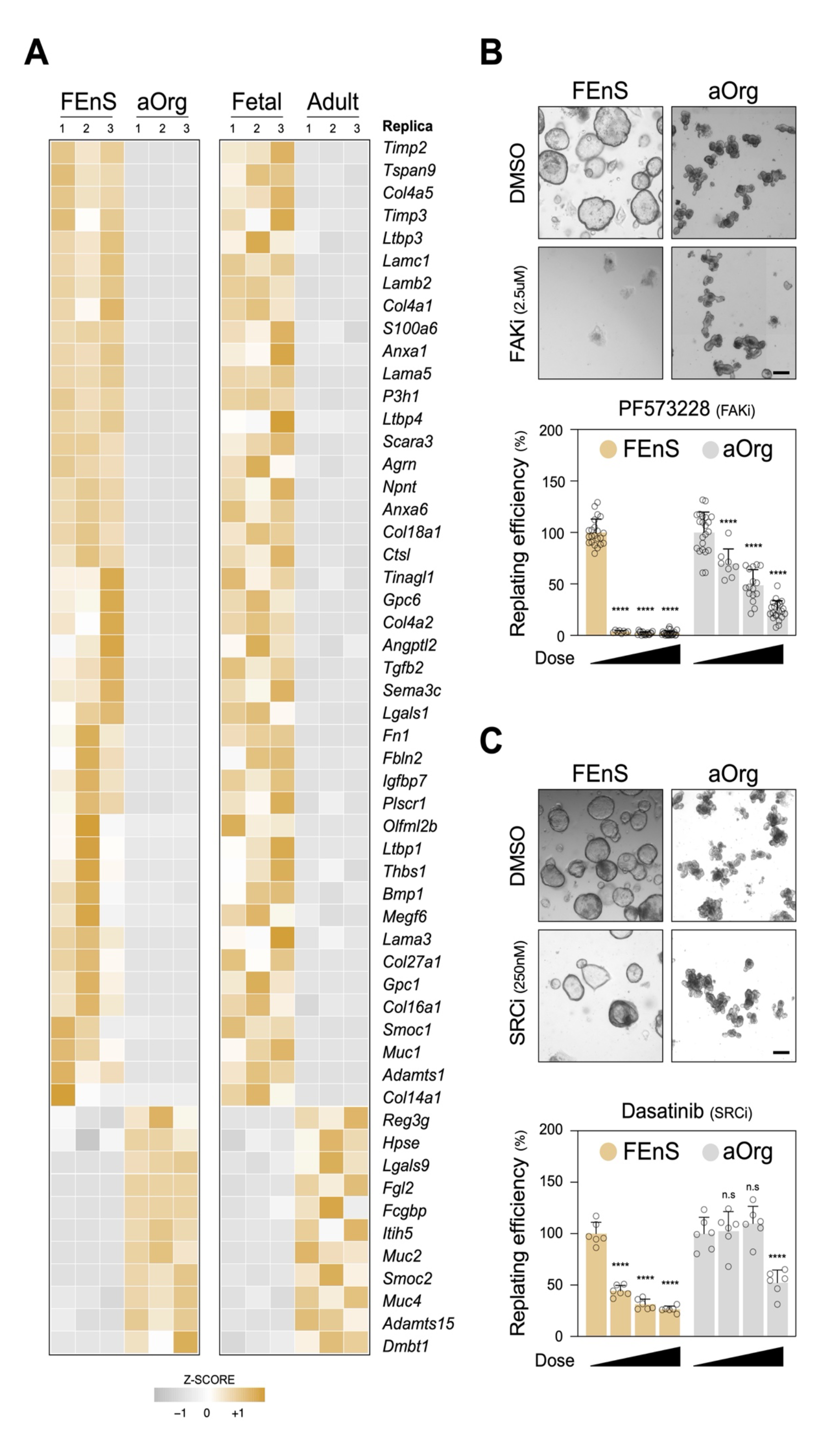
**(A)** Heatmap showing expression of ECM genes (GO: 0031012 - Extracellular matrix) that are differentially expressed consistently between FEnS and aOrg, and freshly isolated fetal and adult epithelial cells. (**B-C)** Replating assay of FEnS and aOrg cultures treated with the pharmacological inhibitors of FAK (B) and SRC (C) kinases. Each dot represents a technical replica of at least three independent experiments. n.s. for non-significant, and **** for p-value < 0.0001 by two-way ANOVA following correction for multiple comparisons (each dose vs 0uM for FEnS or aOrg). Scale bar: 200um.

